# Nucleotide-driven KaiC dynamics coordinate the core properties of the cyanobacterial circadian clock

**DOI:** 10.64898/2026.06.19.733361

**Authors:** Kumiko Ito-Miwa, Tomoaki Muranaka, Takao Kondo, Kazuki Terauchi

**Affiliations:** Department of Earth and Space Science, Graduate School of Science, The University of Osaka, Toyonaka 560-0043, Japan; Institute for Advanced Research, Nagoya University, Nagoya 464-8601, Japan; Department of Biological Science, Graduate School of Science, Nagoya University, Nagoya 464-8602, Japan; College of Life Sciences, Ritsumeikan University, Kusatsu, Shiga 525-8577, Japan; Graduate School of Life Sciences, Ritsumeikan University, Kusatsu, Shiga 525-8577, Japan

**Keywords:** Circadian clock, cyanobacteria, KaiC, ATPase, nucleotide

## Abstract

Circadian clocks generate stable ∼24-h rhythms with a defined period, temperature compensation, and entrainment to external cues that set phase. However, the molecular reactions that generate these features are not fully understood. In cyanobacteria, timekeeping is driven by the hexameric ATPase KaiC, which consists of two homologous domains, CI and CII, and whose enzymatic turnover underlies rhythmic phosphorylation. Here we identify ADP release as the rate-limiting step in the KaiC ATPase cycle and demonstrate that the KaiC ATPase nucleotide cycle integrates core properties of the circadian clock. Across KaiC period mutants, increased ADP occupancy is associated with longer periods. Elevated temperature shifts KaiC toward an ADP-bound state, offsetting the thermal acceleration of ATP hydrolysis. KaiB reinforces this ADP-bound conformation by inhibiting nucleotide exchange, thereby strengthening inhibition of CI ATPase activity of KaiC and tuning oscillation amplitude in a temperature-dependent manner. In contrast, KaiA accelerates ADP-to-ATP exchange within the KaiB–KaiC complex, thereby stimulating CI ATPase activity and promoting KaiB–KaiC dissociation prior to KaiC phosphorylation. Phosphorylation begins only after this transition, indicating that KaiC nucleotide-bound state sets the phase of the phosphorylation cycle. Collectively, these results establish the nucleotide cycle of KaiC ATPase as a unifying mechanism that connects molecular reactions to the defining properties of the cyanobacterial circadian clock.

## Introduction

Circadian clocks synchronize cellular physiology with the Earth’s day–night cycle by generating self-sustained ∼24-h oscillations. Across diverse organisms, circadian systems are characterized by three canonical properties: an approximately 24-h period, entrainment by environmental cues such as light–dark cycles, and temperature compensation, whereby the period remains nearly constant across a range of temperatures (1). Although these defining properties have been extensively characterized at the physiological level, the molecular mechanisms that establish and stabilize them remain incompletely understood. In many organisms, circadian rhythms are explained by transcription–translation feedback loops (TTFLs), in which clock gene products periodically repress their own expression (2). Although TTFL-based models account for oscillatory gene expression, they do not fully explain how circadian systems achieve robust and temperature-compensated timekeeping (3).

Cyanobacteria provide a unique framework for dissecting the molecular mechanisms underlying the defining properties of circadian clocks, including ∼24-h periodicity, entrainment, and temperature compensation (4, 5). A functional circadian oscillator can be reconstituted in vitro using only three proteins—KaiA, KaiB, and KaiC—together with ATP, demonstrating that robust circadian timekeeping can emerge solely from post-translational biochemical reactions (6). Within this system, KaiC serves as the central pacemaker (4). It is a two-domain P-loop ATPase that forms a homohexameric ring composed of an N-terminal CI domain and a C-terminal CII domain (7, 8). ATPase activity associated with period determination resides primarily in CI (9, 10), whereas CII drives the ordered phosphorylation–dephosphorylation cycle at Ser431 and Thr432 (11, 12). Coupling between these domains modulates KaiC interactions with KaiA and KaiB in a phosphorylation-dependent manner, enabling sustained oscillations in vitro (13–15).

KaiC hydrolyzes ATP at an exceptionally low rate (∼10–15 ATP molecules day⁻¹ per hexamer at 30°C), and this ATPase activity is both temperature compensated and tightly correlated with circadian period (Terauchi–Kondo plot) (9). Remarkably, these characteristics persist even in the absence of KaiA and KaiB, suggesting that KaiC itself encodes fundamental timing information. Supporting this view, recent work demonstrated that a KaiC ATPase–driven protein oscillator intrinsically sustains a robust circadian period (16).

Together, these observations suggest that core clock properties are embedded in the structural and enzymatic design of KaiC. To account for this behavior, we previously proposed that KaiC ATPase activity is self-regulated via a structural negative-feedback mechanism (4, 9). Such self-limiting control, implemented through intrinsically slow biochemistry, could provide a molecular basis for both temperature compensation and period determination. ATPase typically undergo catalytic cycles involving ATP binding, hydrolysis, and ADP release—i.e., nucleotide exchange between ATP- and ADP-bound states(17–19); KaiC is no exception. These nucleotide-dependent states are expected to reshape conformational dynamics in CI and CII and, in turn, govern progression through the circadian reaction cycle (20). However, how KaiC nucleotide occupancy-state in KaiC hexamer is mechanistically coupled to ATPase turnover and structural transitions—and how this coupling specifies period length, temperature compensation, and phase—remains unclear. Here we define the biochemical basis of this regulation and show that the nucleotide state of KaiC is a central determinant of the dynamics of the cyanobacterial circadian oscillator.

## Results

### Transient dynamics reveal feedback regulation of KaiC ATPase activity

KaiC ATPase activity is a major determinant of both circadian period and temperature compensation in the cyanobacterial clock system (9, 10, 21). We previously showed that the intrinsic ATPase activity of KaiC, even in the absence of KaiA or KaiB, determines the circadian period in period-altered strains (9). Here we asked whether temperature compensation of KaiC ATPase activity is preserved in these period mutants. The Q_10_ values of ATPase activity for the KaiC mutants S157P, F470Y, T42S, and A251V were close to unity (1.04, 0.94, 0.93, and 0.96, respectively), comparable to that of wild-type KaiC (Fig. 1A). Thus, temperature compensation of KaiC ATPase activity remains largely intact in these mutants, supporting the view that KaiC ATPase activity itself is central to temperature compensation of the circadian clock.

**Figure 1.**
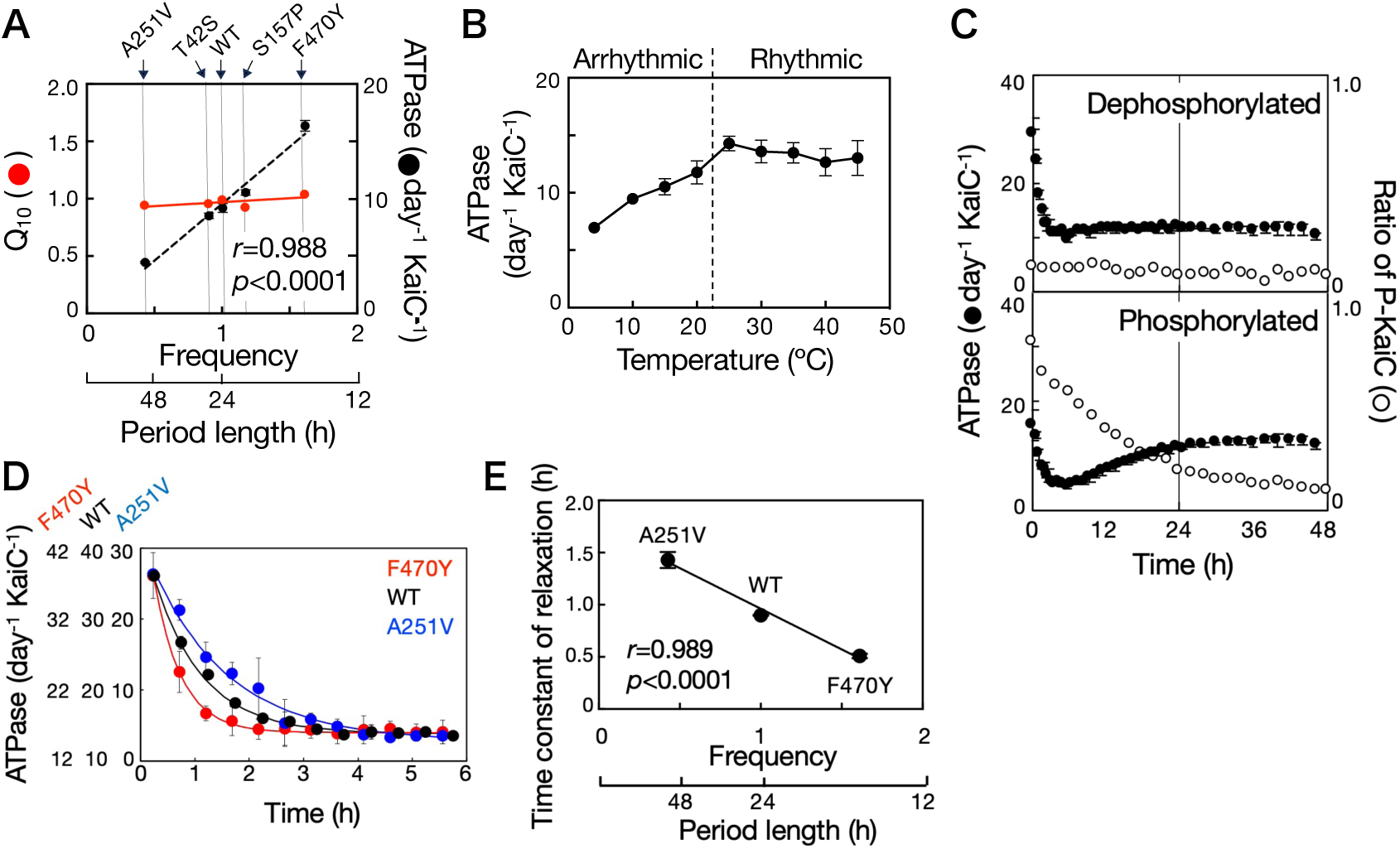
Intramolecular negative feedback regulation of KaiC ATPase activity. (*A*) Temperature compensation of ATPase activity of wild-type (WT) KaiC and period-altered KaiC variants (A251V, T42S, S157P, and F470Y). KaiC proteins were incubated for 2 d at 20, 30, or 40°C in the absence of KaiA and KaiB. ATPase activity (black circles) and the corresponding Q_10_ values (red circles) are plotted as a function of the frequencies of the in vitro phosphorylation cycle. ATPase activity data are presented as mean ± SD (n = 3) (*r*, correlation coefficient; *p*, *P* value). Frequencies of the in vitro phosphorylation cycle were obtained from (16). (*B*) ATPase activity of KaiC in the absence of KaiA and KaiB. KaiC protein was incubated for 2 d at 4, 10, 20, 25, 30, 35, 40, and 45°C. Data are presented as mean ± SD (n = 6 for 30°C; n = 3 for all other temperatures). (*C*) Transient ATPase responses (black circles) of dephosphorylated (top) and phosphorylated (bottom) KaiC. Prior to the measurements, phosphorylated and dephosphorylated KaiC were prepared by incubating KaiC at 4°C and 30°C, respectively, for 2 days. At time 0, the KaiC reaction mixture lacking KaiA and KaiB was shifted from ice to 30°C. Phosphorylation states are overlaid (white circles). Data are presented as mean ± SD for ATPase activity of dephosphorylated KaiC (n = 5) and phosphorylated KaiC (n = 3). Phosphorylation-state data are from a single experiment (n = 1). (*D*) Overlaid transient ATPase trajectories of dephosphorylated KaiC following a temperature step-up from ice to 30°C: WT (black circles), F470Y (red circles), and A251V (blue circles). Curves indicate nonlinear fits. Data are presented as mean ± SD (WT, n = 3; F470Y, n = 4; A251V, n = 5). (*E*) Relationship between ATPase relaxation kinetics and the oscillation frequency of in vitro KaiC phosphorylation rhythm. Time constant of relaxation extracted from the ATPase transients were plotted against the oscillation frequencies of the in vitro phosphorylation rhythms. *r*, correlation coefficient; *p*, *P* value. Frequencies of the in vitro phosphorylation cycle were obtained from (16).

Previous studies have shown that KaiC ATPase activity remains temperature compensated between 25 and 35°C even in the absence of KaiA and KaiB (9). By contrast, KaiC phosphorylation rhythms become damped and ultimately cease below ∼20°C (22). To test whether temperature compensation of ATPase activity persists under conditions where phosphorylation rhythms fail, we measured KaiC ATPase activity over a broader temperature range (4–45°C) (Fig. 1B). KaiC ATPase activity remained temperature compensated between 25 and 45°C (Q_10_ = 0.96), consistent with the temperature range over which phosphorylation rhythms persist. In contrast, temperature compensation was impaired from 4 to 20°C (Q₁₀ = 1.39), suggesting that temperature-compensated ATPase turnover is coupled with sustained robust circadian oscillations.

We previously proposed that conformational transitions driven by the KaiC ATPase cycle generate negative feedback on ATPase activity, thereby enabling temperature compensation by maintaining a low level of enzymatic activity (9). This model predicts that a sudden temperature upshift induces an immediate increase in ATPase activity, followed by a slower attenuation as the negative feedback is established.

To test this prediction, we monitored KaiC ATPase activity following a temperature step-up assay from 4°C to 30°C (Fig. 1C). To keep the CII phosphorylation state constant during the assay, wild-type KaiC (WT-KaiC) was incubated at 30°C for 2 days to be dephosphorylated before experiments (23), WT-KaiC showed an immediate increase in ATPase activity after the temperature-step up; however, this increase was transient, and the activity gradually relaxed to a new steady-state level. By contrast, the well-characterized *E. coli* ATPase RecA (24, 25) exhibited no such transient behavior: its ATPase activity remained elevated after the temperature increase, consistent with a temperature-dependent change in catalytic rate (Fig. S1). This comparison suggests that the transient attenuation is a distinctive property of KaiC.

When phosphorylated KaiC was used in the same assay, ATPase activity showed a pronounced transient decrease with a characteristic undershoot during autodephosphorylation (Fig. 1C). This behavior was not observed in dephosphorylated KaiC or in the catE1^−^ (E77Q/E78Q) mutant (Fig. S2), arguing against the possibility that the undershoot arises from ATP regeneration coupled to dephosphorylation (23, 26). Instead, the undershoot indicates that phosphorylation state modulates KaiC ATPase activity. Together, these findings suggest that KaiC dynamically regulates its ATPase activity via a negative autoregulatory feedback mechanism, consistent with a role in temperature compensation.

To relate these feedback dynamics to circadian timing, we analyzed KaiC variants with altered circadian periods. Using the temperature step-up assay, we measured the transient ATPase response of a short-period mutant (F470Y) and a long-period mutant (A251V), both starting from the dephosphorylated state. After the temperature step-up, F470Y relaxed to the new steady state more quickly than WT-KaiC, whereas A251V relaxed substantially more slowly (Fig. 1D). Notably, the time constant of relaxation closely correlated with the frequency of the phosphorylation cycle (Fig. 1E), indicating that feedback dynamics of KaiC ATPase activity are coupled to the intrinsic frequency of the circadian oscillator.

### ADP release is the rate-limiting step in the KaiC ATPase cycle

To clarify the molecular basis of the negative autoregulation of KaiC ATPase activity observed in Fig. 1, we sought to identify the rate-limiting step in the ATPase cycle. The ATPase cycle generally consists of three steps: ATP binding, ATP hydrolysis (phosphoanhydride bond cleavage), and release of the products ADP and inorganic phosphate (Pi) (17). KaiC ATPase activity is typically quantified by measuring ADP released into solution (9, 16, 21, 27). Because the chemical step of ATP hydrolysis is expected to accelerate with increasing temperature, we reasoned that slow ADP dissociation might instead limit nucleotide turnover in KaiC, potentially explaining its unusually low ATPase activity and weak temperature dependence. To test this idea, we quantified the steady-state nucleotide-bound composition of KaiC by UPLC.

Before incubation, more than 90% of the KaiC-bound nucleotide was ATP when samples were kept on ice (Fig. 2A). After incubation at 30 °C for 48 h, the ATP/total nucleotide ratio decreased to ∼0.3, similar to values reported in previous studies with less than 10-hour incubations (23, 28). indicating that ATP hydrolysis proceeded continuously during incubation. To assess temperature dependence, we incubated KaiC at 25, 30, 35, and 40 °C and quantified bound nucleotides by UPLC (Fig. 2A). The ATP/total nucleotide ratio decreased progressively with increasing temperature, showing that higher temperatures shift KaiC toward the ADP-bound state. Because each KaiC protomer contains two nucleotide-binding sites that are fully occupied by either ATP or ADP (23, 28), this shift reflects temperature-dependent accumulation of protein-bound ADP, consistent with the expected thermal acceleration of the chemical hydrolysis step. In contrast, the rate of ADP release into solution, which is typically used as a measure of KaiC ATPase activity, remained nearly constant from 25 °C to 45 °C (Fig. 1B). Together, these results indicate that ADP release, rather than ATP hydrolysis itself, is the rate-limiting step in the KaiC ATPase cycle. Moreover, ADP/total nucleotide ratios exceeding 0.5 suggest preferential ADP retention, likely in the CI domain, at elevated temperatures.

**Figure 2.**
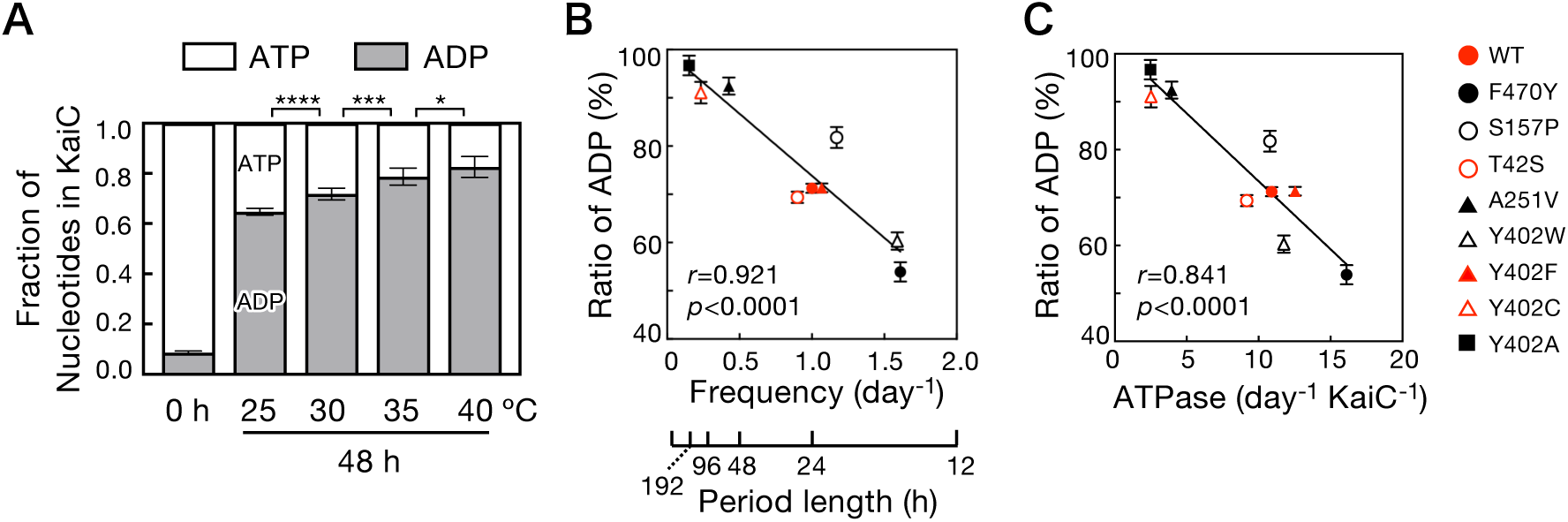
ADP release limits turnover in the KaiC ATPase cycle. (*A*) Temperature dependence of KaiC nucleotide occupancy. KaiC was incubated for 48 h at 25, 30, 35, or 40°C. At each temperature, aliquots were collected, free nucleotides were removed, and the remaining KaiC-bound nucleotides were quantified. The fractions of ATP and ADP relative to the total pool of bound nucleotides are plotted as a function of temperature. Statistics were evaluated by unpaired *t* tests with Welch’s correction; significant differences are indicated (\**P* < 0.05, *** *P* < 0.0005, **** *P* < 0.0001). Data are mean ± SD (n = 6). (*B, C*) Steady-state nucleotide occupancy of WT KaiC and period mutants. The fraction of ADP among total KaiC-bound nucleotides is plotted against the frequency of the in vitro phosphorylation cycle (*B*) or against KaiC ATPase activity (*C*). *r*, correlation coefficient; *p*, *P* value. Nucleotide occupancy data are presented as mean ± SD (WT, n = 5; each mutant, n = 3). Frequencies of the in vitro phosphorylation cycle were obtained from (16, 21).

To evaluate the physiological relevance of ADP retention, we quantified steady-state nucleotide occupancy in KaiC period mutants. KaiC ATPase activity is inversely related to circadian period length, such that lower ATPase activity is associated with longer periods (9, 16, 21). Consistent with this relationship, steady-state ATP/total nucleotide ratios across mutants correlated with both ATPase activity and period length (Figs. 2B, 2C): mutants with higher ADP fractions exhibited lower ATPase rates and longer circadian periods. In long-period mutants (e.g., Y402A, Y402C, and A251V), ADP comprised >90% of the KaiC-bound nucleotide, indicating predominant ADP retention in both the CI and CII domains. Such strong ADP occupancy would be expected to suppress ADP release and thereby slow subsequent nucleotide exchange/ATP loading into CI, reducing overall ATPase turnover. The Y402L mutant was a notable exception, showing only a weak correspondence between nucleotide occupancy and ATPase activity (Fig. S3). Together, these results support ADP release kinetics as a key determinant of KaiC ATPase activity and circadian timing.

### Differential temperature dependence of CI and CII ATPase activities in KaiC

In the presence of KaiA and KaiB, KaiC ATPase activity oscillates in concert with the KaiC phosphorylation cycle (9, 29). However, because KaiC exhibits extremely low ATPase activity, this ATPase activity oscillation has been difficult to track at high temporal resolution. To overcome this limitation, we developed a UPLC-based assay that quantifies KaiC ATPase activity with high temporal resolution at 30 °C (Fig. 3A).

**Figure 3.**
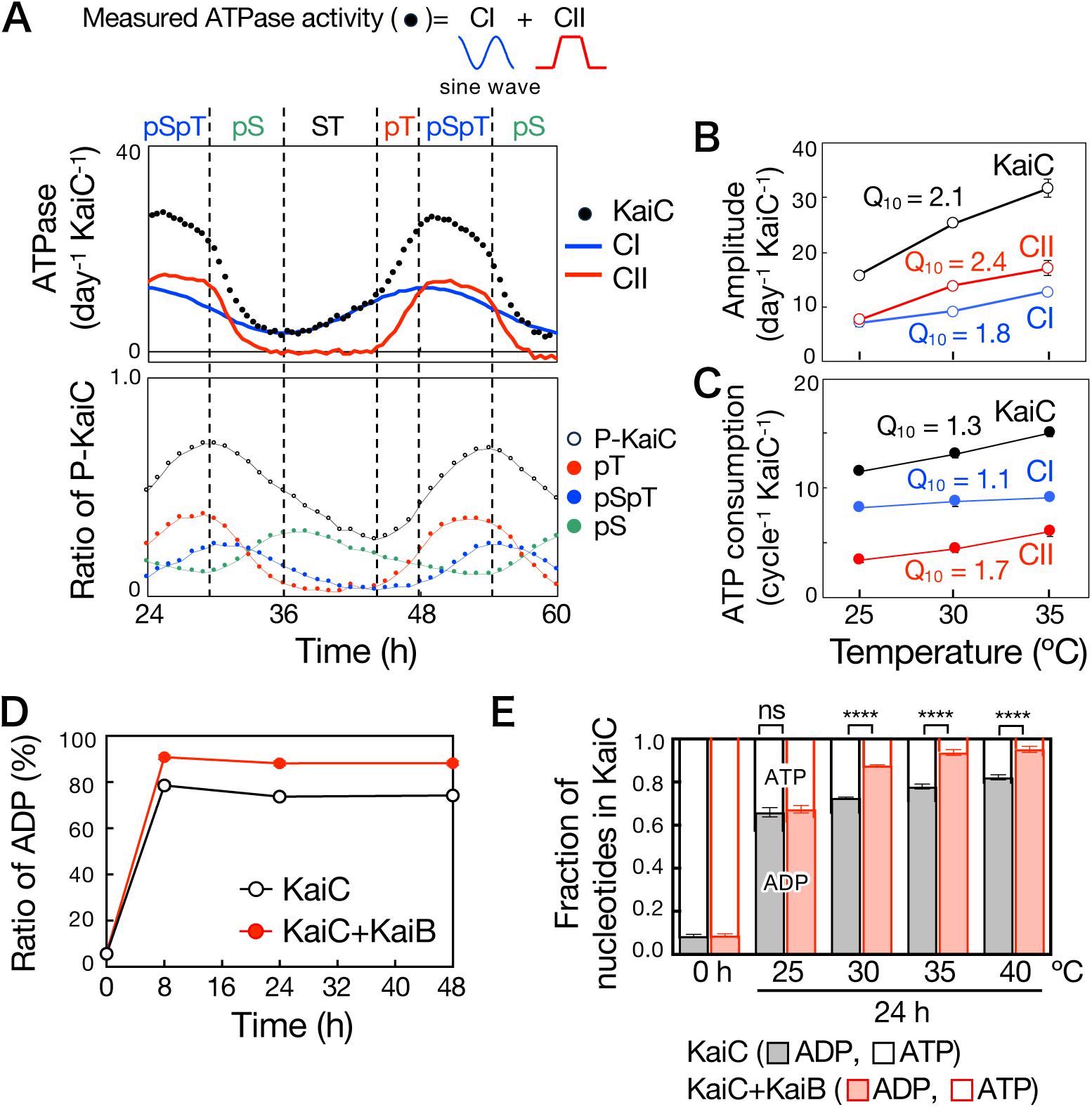
Circadian oscillations of KaiC ATPase activity in the presence of KaiA and KaiB. (A) Circadian oscillations of KaiC ATPase activity (top) and KaiC phosphorylation (bottom) in an in vitro reconstituted KaiABC reaction at 30°C. Black circles denote measured KaiC ATPase activity. Blue and red curves indicate the inferred CI- and CII-derived ATPase components, respectively. Data are presented as mean values (ATPase activity, n = 8; phosphorylation, n = 11). (*B, C*) Oscillation amplitude (peak-to-trough) (*B*) and ATP consumed per cycle (*C*) calculated from the measured total ATPase activity and the inferred CI and CII components. Amplitude was defined as the difference between the maximum and minimum ATPase activities (peak-to-trough) over the oscillation cycle. Data are presented as mean ± SD (n = 3 for 25°C and 35°C; n = 8 for 30°C). (*D*) Effect of KaiB on KaiC nucleotide occupancy at 30°C. The fraction of ADP among total KaiC-bound nucleotides is plotted over time. Data are mean ± SD (n = 3). (*E*) Temperature dependence of KaiC nucleotide occupancy in the presence or absence of KaiB. KaiC with or without KaiB was incubated for 24 h at 25, 30, 35, or 40°C. Statistical significance was assessed using unpaired t tests with Welch’s correction; significance levels are indicated as ns (not significant) and **** (*p* < 0.0001). Data are presented as mean ± SD (n = 4).

With this assay, KaiC ATPase activity displayed a characteristic, reproducible oscillatory waveform that closely tracks phosphorylation at S431 and T432 (11, 12). ATPase activity fell steeply during the pSpT→pS/T transition, rose gradually from pST→S/T, increased sharply during ST→S/pT, and then declined more slowly from SpT back to pSpT (S, S431; T, T432; p, phosphorylated). Similar waveforms were observed at 25 °C and 35 °C (Fig. S4). As reported previously (6), the period of the phosphorylation cycle was temperature compensated over this range and we additionally found that the period of ATPase activity was likewise temperature-compensated. By contrast, oscillation amplitude increased with temperature: the ATPase amplitude showed strong temperature dependence (Q_10_ = 2.1), paralleling the temperature-dependent increase in phosphorylation amplitude (Fig. 3B and Fig. S4). Thus, temperature primarily modulates oscillation amplitude while leaving the period largely unchanged.

To determine how the CI and CII domains individually contribute to the oscillatory ATPase dynamics, we attempted to separate CI- and CII-derived ATPase activities from the total signal (Fig. 3A and Fig. S4; blue and red traces, respectively). Previous studies indicate that CII ATPase is negligible in the dephosphorylated state, whereas KaiA activates CII ATPase via the A-loop (28, 29). Thus, ATPase activity during the dephosphorylation phase should largely reflect CI. We therefore fit the dephosphorylation-phase ATPase trace with a simple sine function as an estimate of the CI component, and defined the CII component as the residual obtained by subtracting this fit from the total signal. The resulting CII waveform peaked near the SpT phosphorylation state—early in the phosphorylation phase—supporting the validity of this decomposition (Fig. 3A and Fig. S4).

Domain-resolved analysis showed that ATPase activities in both CI and CII oscillate, and that the oscillation amplitudes increase with temperature (Fig. 3B). However, the two domains differed markedly in temperature sensitivity. Total ATP consumption per oscillation cycle increased only modestly with temperature (Q_10_ = 1.3), whereas CII ATP consumption per cycle increased substantially (Q_10_ = 1.7). In contrast, CI ATP consumption per cycle remained nearly constant across the temperatures tested (Fig. 3C). Thus, CI and CII exhibit distinct temperature responses in their ATPase dynamics.

To investigate why oscillation amplitude increases with temperature, we asked how temperature affects KaiC nucleotide occupancy and KaiB binding. As shown in Fig. 2A, increasing temperature shifts KaiC toward the ADP-bound state. KaiB is known to preferentially bind the ADP-bound CI conformation, especially when CII is in the pSpT or pST state (13, 14). Thus, the temperature-driven shift in CI nucleotide occupancy should influence KaiB association. Consistent with this, adding KaiB at 30 °C increased the ADP/total nucleotide ratio of KaiC (Fig. 3D), suggesting that KaiB stabilizes the ADP-bound CI state by inhibiting ADP release and the subsequent ATP exchange. With increasing temperature, the ATP-bound fraction decreased further (Fig. 3E) and KaiB–KaiC complex formation increased across 25–35 °C (Fig. S5). Together, these results indicate that temperature-driven ADP accumulation promotes KaiB binding to CI. Stronger KaiB association at higher temperature would more strongly suppress CI ATPase, deepen the troughs of the ATPase oscillation, and thereby increase oscillation amplitude. Notably, because KaiC equilibrates to a dephosphorylated state at 30 °C (Fig. 1C and Fig. S2), our observation that KaiB increases the ADP/total nucleotide ratio indicates that KaiB inhibits CI ADP release and ATP rebinding even when CII is not in the pSpT or pST state.

### CI nucleotide exchange triggers KaiC activation

The nucleotide occupancy state of KaiC governs its biochemical properties throughout the circadian cycle (Figs. 2 and 3). To determine how nucleotide exchange contributes to CII phosphorylation, we tracked the timing of changes in nucleotide binding, ATPase activity, and phosphorylation during the transition from the inactive (dephosphorylation) to the active state (phosphorylation). This transition was triggered by adding KaiA to a preincubated KaiB–KaiC complex and then biochemical changes in KaiC were monitored over time (Fig. 4). To resolve the order of events, we used 0.6 µM KaiA (half the concentration used under standard oscillatory conditions). This subsaturating KaiA slowed the reaction without altering the activation pathway, enabling us to resolve intermediate steps with improved temporal resolution.

**Figure 4.**
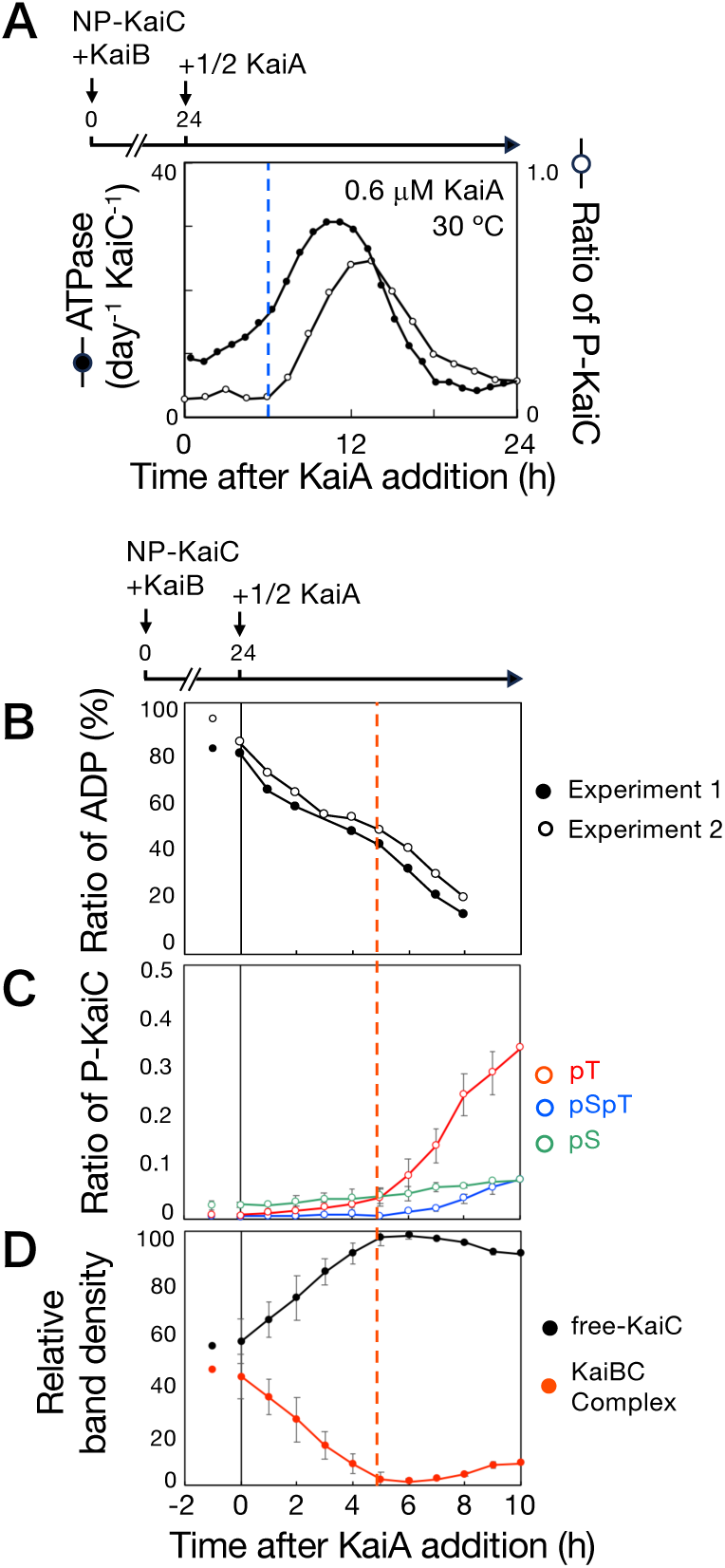
CI-ATPase activation precedes and enables KaiC phosphorylation. *(A)* Time course showing stimulation of KaiC ATPase activity (filled circles) before the onset of KaiC phosphorylation (open circles) after addition of KaiA at half the standard concentration at 30°C. Corresponding results at 35°C are shown in Fig. S7. The blue dashed line marks the initiation of phosphorylation. (*B* –*D*) Time courses of KaiC nucleotide occupancy (*B*), KaiC phosphorylation level (*C*), and KaiB–KaiC complex assembly (*D*) following addition of KaiA at half the standard concentration. Dephosphorylated KaiC was preincubated with KaiB for 24 h at 30°C, after which KaiA (0.6 μM) was added. For nucleotide-occupancy measurements, aliquots were collected at the indicated times, free nucleotides were removed, and bound nucleotides were quantified. For analysis of KaiC phosphorylation and KaiB–KaiC complex formation, aliquots were collected at the indicated times and analyzed by modified BN-PAGE or SDS-PAGE. BN-PAGE bands corresponding to free KaiC hexamers and KaiB–KaiC complexes were quantified by densitometry (Fig. S6), and the fraction of complexed KaiC was calculated as the band intensity of the complex divided by total KaiC signal (complex + free hexamer). The red dashed line indicates the time at which phosphorylation begins and the KaiB–KaiC complex disappears. Data are presented as mean ± SD (phosphorylation, n = 6; KaiB–KaiC complex assembly, n = 3).

Under these conditions, KaiA addition led to an immediate exchange of bound ADP for ATP in KaiC, accompanied by rapid stimulation of ATPase activity (Figs. 4A and 4B). By contrast, KaiC phosphorylation was delayed by ∼5 h (Figs. 4A and 4C). During this lag, KaiB progressively dissociated from KaiC, increasing the fraction of free (unbound) KaiC, and phosphorylation began only after the KaiB–KaiC complex had fully disassembled (Fig. 4D and Fig. S6). Thus, KaiC activation proceeds in a defined sequence: nucleotide exchange and ATPase activation occur first, KaiB release follows, and CII autophosphorylation is initiated only after these preceding steps. Consistently, phosphorylation onset coincided with near-complete loss of the KaiBC complex, indicating that KaiB dissociation precedes—and is required for—initiation of CII autophosphorylation.

In Fig. 4B, the red trace marks the time when the ADP/total nucleotide ratio reaches ∼50%, i,e. ATP/total is also ∼50%. By this time, the KaiBC complex has largely dissociated and phosphorylation begins (Fig. 4D and Fig. S6). Previous studies showed that complete KaiB release from KaiC requires KaiA-dependent activation of CI and CII ATPase activities (29). Because KaiB inhibits ADP dissociation from CI (Fig. 3), KaiB dissociation from CI should restore CI nucleotide exchange and CI ATPase activity. Notably, CII remains unphosphorylated until this transition, suggesting that the CI nucleotide state gates the onset of phosphorylation. An ATP/total ratio of ∼50% may therefore reflect preferential ATP loading in CI, implying that CII phosphorylation starts only after CI becomes predominantly ATP bound.

Together, these results support a model in which CI nucleotide exchange—ADP release followed by ATP binding—serves as the molecular trigger for KaiC activation. This transition stimulates CI ATPase activity and promotes dissociation of KaiB from the KaiBC complex. Because phosphorylation begins only after KaiB release, KaiB binding likely maintains KaiC in an inactive state, whereas KaiB dissociation permits initiation of CII autophosphorylation. In this way, CI nucleotide occupancy functions as a regulatory switch that orchestrates the ordered progression from KaiC inactivation to full activation.

### Intrinsic KaiC dynamics determine the phase of the KaiABC oscillator

Our preceding analyses defined how nucleotide exchange and KaiB dissociation control KaiC activation (Figs. 2–4). We next asked whether these intrinsic biochemical processes also underlie another defining circadian property—phase determination. The phase of KaiC phosphorylation rhythms is known to shift in response to perturbations such as temperature changes or altered ATP/ADP ratios (30, 31). Although KaiA and KaiB are required to generate KaiC oscillations, it remains unresolved whether oscillator phase is imposed by the timing of KaiA–KaiB–KaiC interactions or instead emerges from—and is constrained by—KaiC-intrinsic biochemical dynamics.

To discriminate between these alternatives, we initiated oscillations in vitro by adding KaiA at defined delays to a pre-incubated KaiB–KaiC mixture (Fig. 5). If phase were determined solely by the timing of KaiA binding, then KaiA addition at different times should reset the oscillation phase accordingly. Conversely, if phase were specified primarily by KaiC-intrinsic dynamics, the oscillation phase should be relatively insensitive to the timing of KaiA addition.

**Figure 5.**
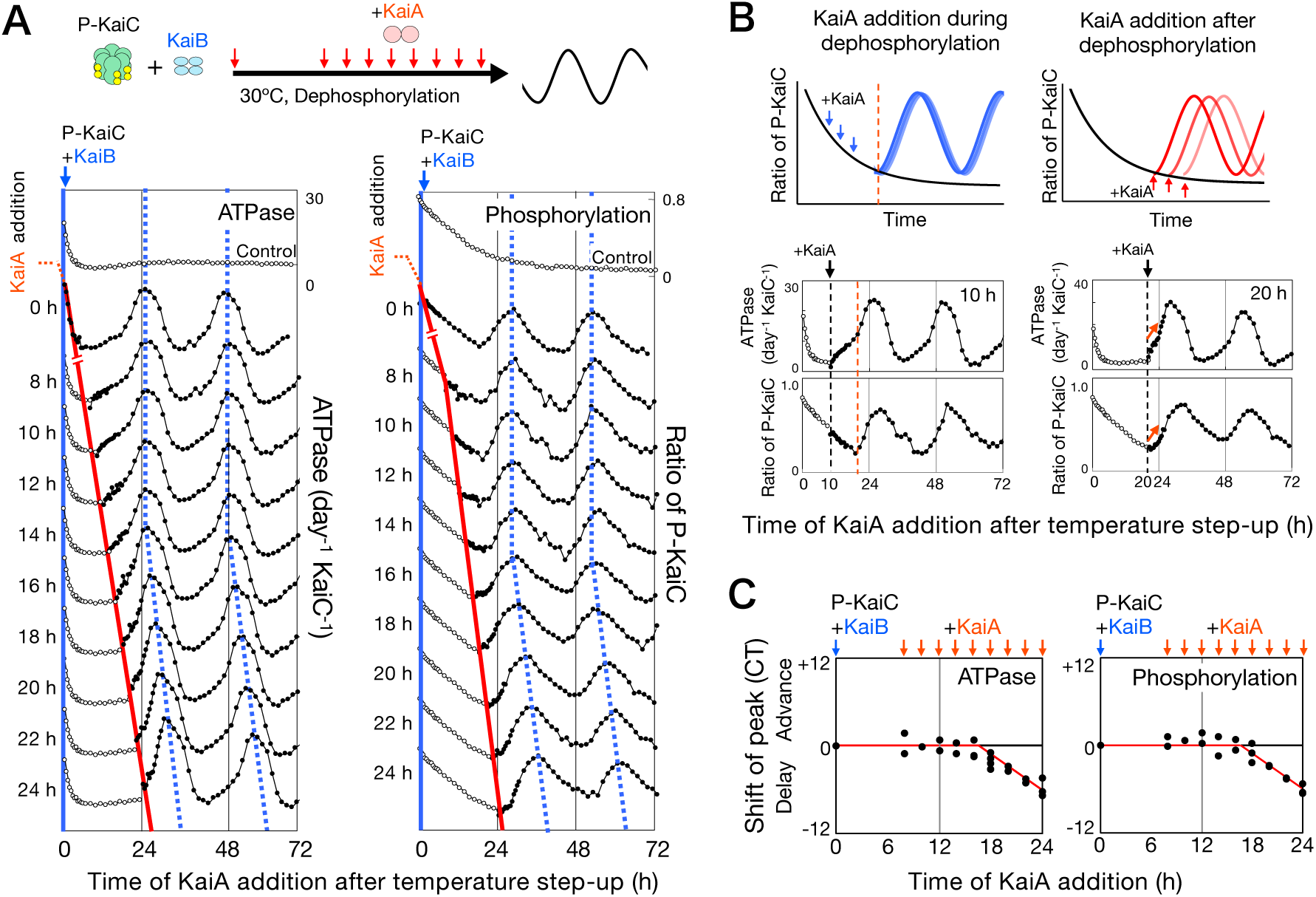
Circadian phase is set by KaiC-intrinsic dynamics. (*A*) Representative time courses of KaiC ATPase activity (left) and phosphorylation state (right). A mixture of phosphorylated KaiC and KaiB was shifted from ice to 30°C at time 0, and KaiA was added at 0, 8, 10, 12, 14, 16, 18, 20, 22, or 24 h after the temperature shift. Open circles (upper traces) and filled circles (lower traces) indicate ATPase activity and phosphorylation state measured in the absence and presence of KaiA, respectively. The red solid line marks the time of KaiA addition. Blue dashed lines indicate the peak positions of the ATPase and phosphorylation rhythms. (*B*) Schematic summary (top): KaiA addition during dephosphorylation caused no detectable phase change (left), whereas addition after dephosphorylation produced time-dependent phase shifts (right). Representative trajectories (bottom) of ATPase activity (upper) and phosphorylation state (lower) following KaiA addition to phosphorylated KaiC in the presence of KaiB at 10 h (left) or 20 h (right). The red dashed line denotes the transition into the phosphorylation phase. (*C*) Phase resetting of KaiC ATPase and phosphorylation rhythms as a function of KaiA addition time. Phase shift was quantified as the difference in the timing of the second peak relative to the control condition (KaiA added at time 0) and plotted against the KaiA addition time. Symbols show individual replicate measurements. Replicate numbers (ATPase/phosphorylation) are: 8 h, 2/2; 10 h, 1/2; 12 h, 2/2; 14 h, 2/2; 16 h, 4/4; 18 h, 4/4; 20 h, 2/2; 24 h, 3/3.

KaiC was pre-incubated with KaiB at 30°C for 8 h, which is sufficient to form a stable KaiBC complex (32). KaiA was then added to separate reactions at 2-h intervals (i.e., at defined delays), and ATPase activity and phosphorylation were monitored after KaiA addition (Fig. 5). KaiA was used at the standard molar ratio for in vitro oscillations, i.e., twice the concentration used in Fig. 4. For KaiA additions up to 16 h, the phase of the phosphorylation rhythm was largely unchanged despite differences in KaiA addition timing (Fig. 5A). By contrast, when KaiA was added at 18 h or later, the phase shifted progressively in a time-dependent manner (Fig. 5A), suggesting a state transition in KaiC around ∼16–18 h that alters its responsiveness to KaiA. Consistent with this, KaiA addition during the dephosphorylation phase (e.g., 10 h) rapidly stimulated ATPase activity while phosphorylation continued to decline (Fig. 5A; Fig. 5B, left). In contrast, when KaiA was added after full dephosphorylation (e.g., 20 h), ATPase activity and phosphorylation increased concurrently (Fig. 5A; Fig. 5B, right). Overall, these data suggest that the phase of KaiC oscillations is not dictated solely by the external timing of KaiA binding (Fig. 5C). Together, they indicate that phase progression is constrained by KaiC-intrinsic dynamics, with KaiC providing an internal timing framework for the KaiABC oscillator.

## Discussion

Although the KaiABC oscillator has been studied extensively, it remains unclear how ATP hydrolysis by KaiC is mechanistically linked to core circadian properties, including period determination, temperature compensation, and phase regulation. Here we identify the nucleotide state of KaiC as a unifying molecular principle underlying these defining features of the cyanobacterial clock. KaiC—the central component of the oscillator—hydrolyzes ATP at an exceptionally low rate, a property required to sustain ∼24-h rhythmicity (9, 10). Our data indicate that the CI-domain nucleotide cycle acts as a regulatory module that governs progression through the oscillator. In this scheme, ATP hydrolysis progressively increases ADP occupancy within CI, and accumulated ADP in turn suppresses further turnover, generating a negative-feedback that stabilizes the ATPase cycle. This nucleotide-dependent regulation buffers ATPase activity against thermal acceleration and modulates KaiC interactions with KaiA and KaiB, providing a mechanistic framework linking ATPase dynamics to circadian period, temperature compensation, and phase in the KaiABC oscillator.

Consistent with the unusually slow ATPase activity of KaiC, our temperature step-up assay reveals an intrinsic feedback architecture within the KaiC ATPase cycle (Fig. 1). After temperature step-up, KaiC ATPase activity exhibits a transient overshoot followed by relaxation to a new steady level (Fig. 1C). This overshoot–recovery signature is a hallmark of negative feedback and is not observed for a non-circadian ATPase such as RecA (Fig. S1), suggesting that the behavior reflects an embedded regulatory property of the KaiC system rather than a simple thermodynamic consequence of temperature on catalysis. Notably, the relaxation timescale covaries with the free-running period across conditions (Figs. 1D and 1E), suggesting that the same internal process that sets circadian timing also governs recovery of ATPase activity after perturbation. Together, these observations are consistent with a post-hydrolysis, negative feedback mechanism that transiently restrains catalytic turnover and stabilizes the ATPase cycle.

Our initial observations (Fig. 1) raised the question of what molecular feature generates this intrinsic negative feedback. The results in Figs. 2–4 support a model in which the feedback arises from nucleotide processing itself. ATP hydrolysis generates ADP that remains bound within the CI domain for extended periods, and this persistent product occupancy suppresses subsequent ATP turnover by restricting nucleotide exchange. Thus, the reaction product acts as an internal inhibitor, establishing a negative-feedback in KaiC.

A mechanistic basis for this negative feedback can be provided by the nucleotide exchange step (Fig. S8). Following hydrolysis, ADP must dissociate before ATP can rebind, creating a potential kinetic bottleneck. Our measurements indicate that nucleotide exchange is slower than the hydrolysis step and covaries with oscillator period (Fig. 2), consistent with ADP release being rate limiting. This conclusion is further supported by the observation that elevated temperature accelerates ATP hydrolysis yet increases ADP accumulation within KaiC (Fig. 2A), consistent with a metastable ADP-bound CI state in which suppressed exchange feeds back to limit further turnover. A conceptually similar product-binding mechanism has been reported for mammalian CKIδ, a core clock kinase implicated in circadian temperature compensation, where increasing temperature weakens substrate binding to the CKIδ–ATP complex but strengthens product (ADP) binding to the CKIδ–ADP complex, providing a mechanistic basis for temperature compensation (33).

Structural considerations offer a plausible explanation for slow nucleotide exchange. In the CI ring, nucleotide binding is stronger than in CII: ATP in CI forms additional intersubunit hydrogen-bonding interactions with the nucleobase that are absent in CII (7), consistent with tighter nucleotide binding in CI (34). Stronger binding would be expected to slow nucleotide dissociation, implying that ADP release is particularly hindered in CI. Moreover, because nucleotide-binding pockets reside at subunit interfaces within the KaiC hexamer (7), ADP release likely requires cooperative rearrangements of the ring. These constraints provide a structural rationale for slow nucleotide exchange and support the view that ADP release from the CI domain is the rate-limiting transition in the KaiC ATPase cycle.

By strongly constraining the ADP-bound inactive–to–ATP-bound active transition, this mechanism renders KaiC ATPase kinetics resistant to temperature-dependent acceleration and thereby supports temperature compensation of period. Consistent with this interpretation, long-period KaiC variants display higher steady-state ADP occupancy than short-period variants (Fig. 2B), implicating the ADP dissociation rate as a principal determinant of oscillator speed. Slow ADP release from the CI domain also provides a plausible basis for circadian timescales: because nucleotide exchange is far slower than the underlying chemical hydrolysis step, extended residence in the ADP-bound state effectively dilates the catalytic cycle to hours, yielding ∼24-h periodicity. In this view, reduced ADP off-rate lengthens the inactive dwell of KaiC, delays ATP-dependent reactivation, and consequently extends the circadian period.

High time-resolution ATPase measurements provide insight into how distinct oscillator functions may be attributed to the two KaiC domains (Fig. 3). The ATPase waveform closely tracks the phosphorylation cycle, indicating tight coupling between nucleotide turnover and phosphorylation state (Fig. 3A). Because CII ATPase is negligible in the dephosphorylated state, we estimated the CI contribution by fitting the dephosphorylation-phase ATPase trace with a sine function. This decomposition implies that CI ATPase is higher during phosphorylation than during dephosphorylation, consistent with KaiA-dependent stimulation of ATPase in both CI and CII and with KaiA-driven autophosphorylation (29), although the inferred CI waveform (approximated by the sine fit) remains to be experimentally validated. Structural studies also provide a mechanistic basis for phosphorylation-state–dependent modulation of CI ATPase, in which phosphorylation state shifts the lytic water position in the CI active site (29). Partitioning the total signal reveals divergent temperature responses: CI ATP consumption remains nearly invariant, whereas CII ATP hydrolysis increases with temperature (Fig. 3C). Consistent with this framework, increasing temperature drives KaiC toward higher ADP occupancy and enhances KaiB association (Figs. 3D and 3E), effects expected to preferentially suppress CI ATPase activity and deepen the troughs of the oscillation. Thus, our results suggest that period stability is maintained by CI nucleotide state, whereas KaiB-dependent interactions contribute to the amplitude and shape of the ATPase rhythm. These observations further imply that CI nucleotide dynamics may time KaiB binding and release, thereby coordinating the ordered sequence of ATPase changes, KaiB association, and phosphorylation transitions analyzed below (Fig. 4).

The measurements in Fig. 4 define the temporal order of molecular events accompanying KaiC activation. Upon addition of KaiA to a preformed KaiB–KaiC complex, KaiC rapidly exchanged nucleotides, with bound ADP promptly replaced by ATP and ATPase activity increasing immediately (Figs. 4A and 4B). In contrast, KaiC phosphorylation initiated only after a multi-hour delay and commenced only once the KaiBC complex had largely dissociated. Thus, the inactive-to-active transition follows a defined order: CI nucleotide exchange and ATPase activation precede KaiB dissociation, and CII autophosphorylation begins only thereafter. Notably, ATPase activity exhibited a further increase, reaching its higher level only after the KaiB–KaiC complex had largely disassembled (Fig. 4D and Fig. S6), consistent with relief of KaiB-mediated suppression of CI nucleotide exchange together with additional KaiA-dependent stimulation of CI/CII ATPase through A-loop-mediated conformational change (29). This ordering indicates that CI nucleotide status gates activation. Phosphorylation onset coincided with an ATP-bound fraction of roughly half the total nucleotide pool, consistent with preferential ATP replacement in CI occurring before entry into the phosphorylating state. Because KaiB binding suppresses CI nucleotide exchange and ATPase activity, KaiA-driven KaiB release across the KaiC population appears to be a prerequisite for transition into the CII phosphorylation phase (Fig. S9). Accordingly, the CI nucleotide cycle functions as a regulatory switch that synchronizes ATPase activation, KaiB association, and CII phosphorylation.

These experiments further indicate that oscillator phase is specified primarily by the biochemical state of CI (Fig. 5). When KaiA was introduced at different times following KaiB pre-incubation, ATPase and phosphorylation rhythms were robustly re-established across a wide time window. However, the two outputs did not re-emerge synchronously: phosphorylation rhythms reproducibly lagged ATPase rhythms by several hours, effectively tracking the ATPase cycle. This convergence argues that phase is not dictated by the timing of KaiA addition but is instead stored in an internal state of KaiC. In line with the ordered transitions resolved in Fig. 4, the phase-preserving internal variable corresponds to CI nucleotide occupancy. Because CI nucleotide exchange and ATPase activation occur before KaiB dissociation, and because CII phosphorylation begins only after KaiB release, progression through the cycle is constrained by the CI nucleotide state (Fig. S9). Thus, the timing of ADP release and ATP rebinding in CI determines when KaiB can dissociate and when CII phosphorylation can initiate, thereby setting oscillator phase. In this framework, KaiA is not a classical phase-resetting cue; rather, it serves as a permissive factor that enables progression only once CI has reached the appropriate nucleotide state. Phase is therefore encoded as an intrinsic biochemical variable within KaiC, defined by the CI nucleotide state and its coupling to KaiB association–dissociation dynamics. This coupling directly links nucleotide-exchange kinetics to the temporal organization of the circadian cycle.

Recent reports that RUVBL2 hydrolyzes ∼13 ATP per day and that its ATPase activity scales with circadian period (35, 36) echo the “slow ATPase” design principle established for KaiC, suggesting a conserved period-setting element in eukaryotic clocks. Physical interactions between RUVBL2 orthologs and core clock factors, together with concordant phenotypes in human, Drosophila, and Neurospora, further support a two-tier architecture in which ATPase-driven post-translational pacemakers are coupled to TTFLs. A parallel logic operates in cyanobacteria, where the KaiC post-translational oscillator functions alongside a TTFL that drives rhythmic gene expression (37–39). Across these systems, nucleotide-dependent biochemical cycles appear to serve as core timing modules that generate intrinsically slow dynamics, whereas TTFL networks amplify, disseminate, and coordinate temporal information at the cellular scale. Together with emerging evidence on RUVBL2, our results on KaiC suggest that tuning nucleotide-processing kinetics represents a general design principle for circadian timing across biological systems.

## Materials and Methods

### Preparation of Kai proteins

Recombinant KaiA, KaiB, KaiC, and KaiC mutants were expressed in *Escherichia coli* and purified as reported previously (27), with details as follows. pGEX-6P-1 plasmids harboring individually cloned ORFs of *kaiA*, *kaiB*, or *kaiC* from *Synechococcus elongatus* PCC 7942 were transformed into *E. coli* BL21 for expression as GST-fusion proteins. Cells expressing GST-fusion proteins were harvested, resuspended in protein-specific extraction buffers [20 mM Tris-HCl (pH 8.0), 150 mM NaCl, and 0.5 mM EDTA for KaiA; 20 mM Tris-HCl (pH 8.0), 10 mM NaCl, and 0.5 mM EDTA for KaiB; and 20 mM Tris-HCl (pH 8.0), 150 mM NaCl, 0.5 mM EDTA, 1 mM DTT, 1 mM ATP, and 5 mM MgCl₂ for KaiC], and lysed by sonication. Lysates were clarified by centrifugation (24,000 × *g*), and the resulting supernatants were subjected to affinity purification using glutathione Sepharose 4B (GE Healthcare).

For KaiA and KaiC, GST tags were removed by on-column digestion with PreScission protease (GE Healthcare). For KaiB, GST–KaiB was eluted with buffer containing 50 mM Tris-HCl (pH 8.0), 10 mM NaCl, and 20 mM reduced glutathione, desalted using a HiPrep Desalting 26/10 column (Bio-Rad) to remove glutathione, and subsequently digested with PreScission protease.

KaiA was further purified by sequential Resource Q anion-exchange chromatography (90–450 mM NaCl gradient), hydroxyapatite chromatography using a Bio-Scale CHT2-I column (Bio-Rad), and size-exclusion chromatography using a Superdex 200 column (GE Healthcare). KaiB was further purified by Resource Q anion-exchange chromatography (0–300 mM NaCl gradient), followed by hydroxyapatite chromatography using a Bio-Scale CHT2-I column (Bio-Rad), and size-exclusion chromatography using a Superdex 75 column (GE Healthcare). KaiC was further purified by size-exclusion chromatography using a HiPrep Sephacryl S300 column (GE Healthcare), followed by Resource Q anion-exchange chromatography (90–450 mM NaCl gradient).

### Reconstitution of the in vitro KaiC phosphorylation rhythm

In vitro KaiC phosphorylation rhythms were reconstituted essentially as described previously (27), using reaction buffer composed of 20 mM Tris (pH 7.8), 150 mM NaCl, and 5 mM MgCl₂ supplemented with 1 mM ATP. Unless otherwise indicated, KaiA, KaiB, and KaiC were used at concentrations of 1.2, 3.5, and 3.5 μM, respectively; these conditions are referred to as the standard conditions in this study. The concentration of KaiC was determined using the Quick Start™ Bradford Protein Assay (Bio-Rad) prior to preparation of the reaction mixtures. For reconstitution of phosphorylation rhythms using the KaiC A251V variant, the concentration of KaiA was increased to 2.4 μM. Period and amplitude were calculated as described previously (40).

### ATPase assay using a UPLC system

KaiC ATPase activity was quantified using an ACQUITY UPLC system (Waters) essentially as described (27). ADP were separated from ATP using a BEH C18 column (2.1 × 50 mm, 1.7 μm; Waters) at a flow rate of 0.8 mL min⁻¹ with a mobile phase consisting of 14 mM ammonium phosphate, 7 mM tetrabutylammonium hydrogen sulfate (pH 8.5), and 15% (v/v) acetonitrile. ADP concentrations were determined from the corresponding peak areas. To minimize injection-volume variability, signals were normalized to the sum of ATP and ADP peak areas.

### Measurement of transient ATPase activity responses using a UPLC system

For ice-to-30°C temperature-step up experiments, dephosphorylated and phosphorylated KaiC were prepared by incubating KaiC (1–2 mg mL^−1^) at 30°C and 4°C, respectively, for 2 days (23). In both cases, samples were buffer-exchanged at 4°C through Micro Bio-Spin™ P-30 Gel columns (Bio-Rad) equilibrated with the reaction buffer to remove ADP accumulated during incubation. Buffer exchange was completed within 2 h, during which dephosphorylated KaiC underwent negligible rephosphorylation at 4°C (23).

After quantification using the Quick Start™ Bradford Protein Assay (Bio-Rad), reaction mixtures were prepared on ice at 3.5 μM KaiC in reaction buffer. The temperature shift was applied by transferring tubes to a thermal-block incubator (BI-515, ASTEC) equilibrated at 30°C. After 2.5 min, samples were placed in the ACQUITY UPLC system and the run was initiated. Completion of the temperature jump within 2 min in the thermal block was verified. For time-course measurements, the autosampler was programmed to inject 2 μL every 30 min. *E. coli* RecA was purchased from BioAcademia.

### Nonlinear regression analysis of transient ATPase responses

Data shown in Fig. 1D were individually fitted by nonlinear regression using GraphPad Prism (GraphPad Software) with the exponential decay model:

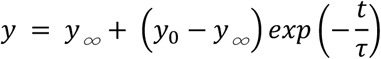

where *y* is KaiC ATPase activity, *y₀* is the KaiC ATPase activity at *t* = 0, *y_∞_* is the steady-state KaiC ATPase activity, *t* is time (h), and *τ* is the time constant of relaxation.

### Detection of KaiC-bound nucleotides

KaiC-bound nucleotides were measured with modifications of a published procedure (28). To evaluate temperature- or mutation-dependent nucleotide occupancy, KaiC hexamers (17.5 μM) were incubated in reaction buffer containing 2 mM ATP. To assess KaiB effects, KaiC and KaiB were mixed at 17.5 μM and 8.8 μM, respectively. Aliquots (70 μL) were collected at the indicated time points and passed once through Bio-Spin P-30 columns equilibrated with ATP-free reaction buffer to remove unbound nucleotides. SDS was added to the eluate (35 μL) to a final concentration of 0.2% (w/v) to release KaiC-bound nucleotides. ATP and ADP were then separated on a BEH C18 column (2.1 × 50 mm, 1.7 μm) using the ACQUITY UPLC system. To confirm ∼2 nucleotides per KaiC protomer, KaiC concentration was determined by the Bradford assay using the remaining eluate prior to SDS addition.

### Estimation of CI- and CII-ATPase activities by phase-specific fitting

Total ATPase activity was modeled as the sum of CI- and CII-derived components. Because KaiA activates CII ATPase activity that drives KaiC phosphorylation, ATPase measurements during the dephosphorylation phase were used to estimate the CI component by fitting a sine function using least-squares fitting in R:

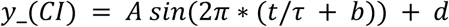

where *y_*(*CI*) is the CI-derived ATPase activity, A is amplitude, τ is period, b is the normalized phase offset (0–1), and d is the mean CI activity. The CII-derived ATPase activity was then obtained as the residual (total minus fitted CI component). CI and CII activities were constrained to be non-negative throughout the oscillatory cycle.

### Blue Native Polyacrylamide Gel Electrophoresis (BN-PAGE)

BN-PAGE was used to assess formation of the KaiB–KaiC complex. BN-PAGE gels were run using a NativePAGE™ Novex Bis-Tris Gel System (Thermo Fisher Scientific) essentially as described (32) with minor modifications. To evaluate temperature effects on complex formation, KaiC (8.8 μM) and KaiB (8.8 μM) were incubated in reaction buffer containing 1 mM ATP. To test the effect of KaiA, KaiA, KaiB, and KaiC were mixed at 1.2 μM (standard) or 0.6 μM (half), 3.5 μM, and 3.5 μM, respectively. For sample preparation, reaction mixtures were combined with 8 μL of NativePAGE sample buffer (Thermo Fisher Scientific) to yield final concentrations of 1× NativePAGE sample buffer, 1 mM ATP, and 5 mM MgCl₂.Because CBB G-250 dissociates KaiC hexamers into gs-KaiC and cs-KaiC (32), it was omitted from the sample buffer to visualize KaiC hexamers as a single band. Samples were separated on NativePAGE™ 4–16% Bis-Tris gels (Thermo Fisher Scientific). Anode and cathode buffers consisted of NativePAGE running buffer containing 5 mM MgCl₂ and 0.25 mM EDTA; the cathode buffer was additionally supplemented with 0.02% CBB and 1 mM ATP, as described previously. Band intensities were quantified by densitometry to determine the relative amounts of KaiB–KaiC complex and free KaiC hexamer.

## Acknowledgments

We gratefully acknowledge our late co-author, Professor Takao Kondo, whose vision, ideas, and passion continue to inspire our work. We also thank Taeko Nishiwaki-Ohkawa and Naoki Takai for their valuable and insightful discussions, and Hisayo Kondo, Tomoe Nishikawa, and Yi Sunyung for their technical assistance. This work was supported by Ohsumi Frontier Science Foundation and NINS Astrobiology Center program research (Grant Number AB0811) to K.I.-M. and by the Japan Society for the Promotion of Science through Grants-in-Aid for Scientific Research (25H02446 and 26K02027 to K.I.-M., 24H02304 to T. M., 17H01427 to T.K., and 19K05833, 24H02301, and 24K01686 to K.T.).

## References

1. C. H. Johnson, “Fundamental propeties of Circadian Rhythms” in Chronobiology Biological Timekeeping, J. J. L. Jay C Dunlap, Patricia J DeCoursey, Ed. (Sinauer Associates, 2004), pp. 67–105.

2. M. W. Young, S. A. Kay, Time zones: a comparative genetics of circadian clocks. Nat Rev Genet 2, 702–715 (2001).

3. T. Kondo, “Around the circadian clock: Review and Preview” in Circadian Rhythms in Bacteria and Microbiomes, C. H. J. a. M. J. Rust, Ed. (Springer, 2021), pp. 21–52.

4. K. Ito-Miwa, K. Terauchi, T. Kondo, “Mechanism of the cyanobacterial circadian clock protein KaiC to measure 24 hours” in Circadian Rhythms in Bacteria and Microbiomes, C. H. Johnson, M. Rust, Eds. (Springer International Publishing,, 2021), pp. 79–91.

5. M. Fang, C. L. Partch, A. LiWang, S. S. Golden, Prokaryotic Circadian Systems: Cyanobacteria and Beyond. Annu Rev Microbiol 79, 523–545 (2025).

6. M. Nakajima et al., Reconstitution of circadian oscillation of cyanobacterial KaiC phosphorylation in vitro. Science 308, 414–415 (2005).

7. R. Pattanayek et al., Visualizing a circadian clock protein: crystal structure of KaiC and functional insights. Mol Cell 15, 375–388 (2004).

8. H. Iwasaki, Y. Taniguchi, M. Ishiura, T. Kondo, Physical interactions among circadian clock proteins KaiA, KaiB and KaiC in cyanobacteria. EMBO J 18, 1137–1145 (1999).

9. K. Terauchi et al., ATPase activity of KaiC determines the basic timing for circadian clock of cyanobacteria. Proc Natl Acad Sci U S A 104, 16377–16381 (2007).

10. J. Abe et al., Circadian rhythms. Atomic-scale origins of slowness in the cyanobacterial circadian clock. Science 349, 312–316 (2015).

11. T. Nishiwaki et al., A sequential program of dual phosphorylation of KaiC as a basis for circadian rhythm in cyanobacteria. EMBO J 26, 4029–4037 (2007).

12. M. J. Rust, J. S. Markson, W. S. Lane, D. S. Fisher, E. K. O’Shea, Ordered phosphorylation governs oscillation of a three-protein circadian clock. Science 318, 809–812 (2007).

13. J. Snijder et al., Structures of the cyanobacterial circadian oscillator frozen in a fully assembled state. Science 355, 1181–1184 (2017).

14. R. Tseng et al., Structural basis of the day-night transition in a bacterial circadian clock. Science 355, 1174–1180 (2017).

15. Y. G. Chang, A. LiWang, The cyanobacterial circadian clock. NPJ Biol Timing Sleep 2, 26 (2025).

16. K. Ito-Miwa, K. Imai, K. Terauchi, T. Kondo, Intrinsic period stability of the cyanobacterial circadian oscillator across in vitro and in vivo conditions. Proc Natl Acad Sci U S A 123, e2526714123 (2026).

17. D. D. Hackney, Kinesin ATPase: rate-limiting ADP release. Proc Natl Acad Sci U S A 85, 6314–6318 (1988).

18. P. I. Hanson, S. W. Whiteheart, AAA+ proteins: have engine, will work. Nat Rev Mol Cell Biol 6, 519–529 (2005).

19. J. P. Erzberger, J. M. Berger, Evolutionary relationships and structural mechanisms of AAA+ proteins. Annu Rev Biophys Biomol Struct 35, 93–114 (2006).

20. Y. Furuike et al., Elucidation of master allostery essential for circadian clock oscillation in cyanobacteria. Sci Adv 8, eabm8990 (2022).

21. K. Ito-Miwa, Y. Furuike, S. Akiyama, T. Kondo, Tuning the circadian period of cyanobacteria up to 6.6 days by the single amino acid substitutions in KaiC. Proc Natl Acad Sci U S A 117, 20926–20931 (2020).

22. Y. Murayama et al., Low temperature nullifies the circadian clock in cyanobacteria through Hopf bifurcation. Proc Natl Acad Sci U S A 114, 5641–5646 (2017).

23. T. Nishiwaki, T. Kondo, Circadian autodephosphorylation of cyanobacterial clock protein KaiC occurs via formation of ATP as intermediate. J Biol Chem 287, 18030–18035 (2012).

24. R. M. Story, T. A. Steitz, Structure of the recA protein-ADP complex. Nature 355, 374–376 (1992).

25. C. E. Bell, Structure and mechanism of Escherichia coli RecA ATPase. Mol Microbiol 58, 358–366 (2005).

26. M. Egli et al., Dephosphorylation of the core clock protein KaiC in the cyanobacterial KaiABC circadian oscillator proceeds via an ATP synthase mechanism. Biochemistry 51, 1547–1558 (2012).

27. K. Ito-Miwa, Y. Onoue, T. Kondo, K. Terauchi, Effect of pH on the cyanobacterial circadian oscillator in vitro. Commun Biol 8, 828 (2025).

28. T. Nishiwaki-Ohkawa, Y. Kitayama, E. Ochiai, T. Kondo, Exchange of ADP with ATP in the CII ATPase domain promotes autophosphorylation of cyanobacterial clock protein KaiC. Proc Natl Acad Sci U S A 111, 4455–4460 (2014).

29. Y. Furuike et al., Regulation mechanisms of the dual ATPase in KaiC. Proc Natl Acad Sci U S A 119, e2119627119 (2022).

30. T. Yoshida, Y. Murayama, H. Ito, H. Kageyama, T. Kondo, Nonparametric entrainment of the in vitro circadian phosphorylation rhythm of cyanobacterial KaiC by temperature cycle. Proc Natl Acad Sci U S A 106, 1648–1653 (2009).

31. M. J. Rust, S. S. Golden, E. K. O’Shea, Light-driven changes in energy metabolism directly entrain the cyanobacterial circadian oscillator. Science 331, 220–223 (2011).

32. K. Oyama, C. Azai, K. Nakamura, S. Tanaka, K. Terauchi, Conversion between two conformational states of KaiC is induced by ATP hydrolysis as a trigger for cyanobacterial circadian oscillation. Scientific Reports 6 (2016).

33. Y. Shinohara et al., Temperature-Sensitive Substrate and Product Binding Underlie Temperature-Compensated Phosphorylation in the Clock. Mol Cell 67, 783–798.e720 (2017).

34. F. Hayashi et al., Roles of two ATPase-motif-containing domains in cyanobacterial circadian clock protein KaiC. J Biol Chem 279, 52331–52337 (2004).

35. M. Liao et al., The P-loop NTPase RUVBL2 is a conserved clock component across eukaryotes. Nature 642, 165–173 (2025).

36. Y. Liu, R. Huo, E. E. Zhang, Evolving perspectives on the molecular and neural foundations of mammalian circadian rhythms. Trends Neurosci 48, 904–918 (2025).

37. S. E. Cohen, S. S. Golden, Circadian Rhythms in Cyanobacteria. Microbiol Mol Biol Rev 79, 373–385 (2015).

38. Y. Kitayama, T. Nishiwaki, K. Terauchi, T. Kondo, Dual KaiC-based oscillations constitute the circadian system of cyanobacteria. Genes Dev 22, 1513–1521 (2008).

39. S. W. Teng, S. Mukherji, J. R. Moffitt, S. de Buyl, E. K. O’Shea, Robust circadian oscillations in growing cyanobacteria require transcriptional feedback. Science 340, 737–740 (2013).

40. M. Nakajima, H. Ito, T. Kondo, In vitro regulation of circadian phosphorylation rhythm of cyanobacterial clock protein KaiC by KaiA and KaiB. FEBS Lett 584, 898–902 (2010).

