## Supporting Information for "Nucleotide-driven KaiC dynamics coordinate the core properties of the cyanobacterial circadian clock"

**This PDF file includes:**

Figures S1 to S9
SI References

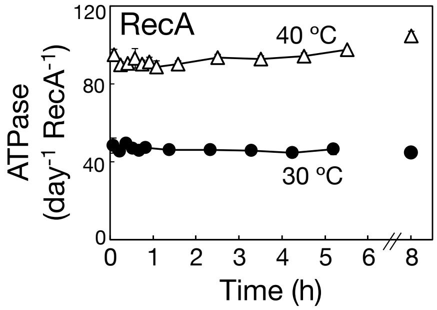


**Fig. S1. Responses of *E. coli* RecA ATPase activity following a temperature step-up from ice to 30°C (black circles) or 40°C (open triangles).**

ATPase activity remained elevated, showing no transient relaxation. Data are presented as mean ± SD (n = 3).

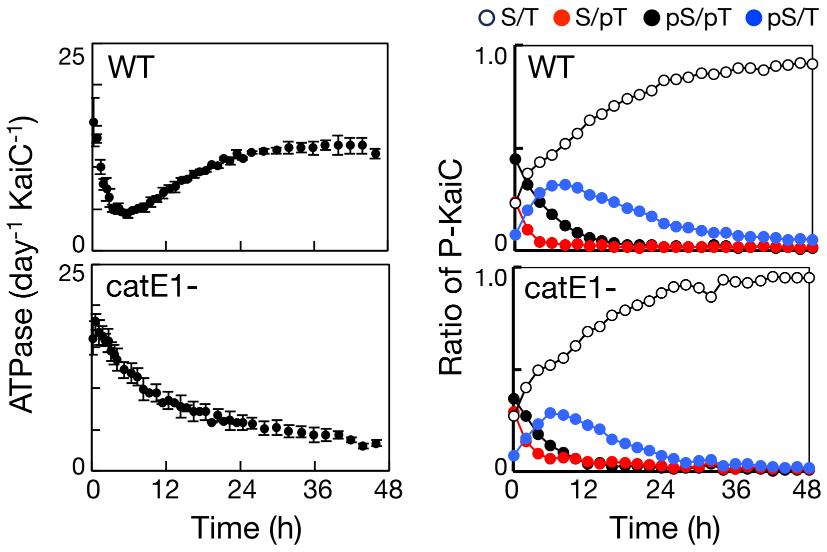


**Fig. S2. Transient responses of ATPase activity (left) and phosphorylation state (right) of WT KaiC and the catE1$^{-}$ (E77Q/E78Q) mutant after an upward temperature shift.**

KaiC was prephosphorylated by incubation at 4°C for 2 d, and the reaction mixture lacking KaiA and KaiB was rapidly transferred from ice to 30°C at time 0. Data are presented as mean ± SD for ATPase activity (n = 3). Phosphorylation-state data are from a single experiment (n = 1).

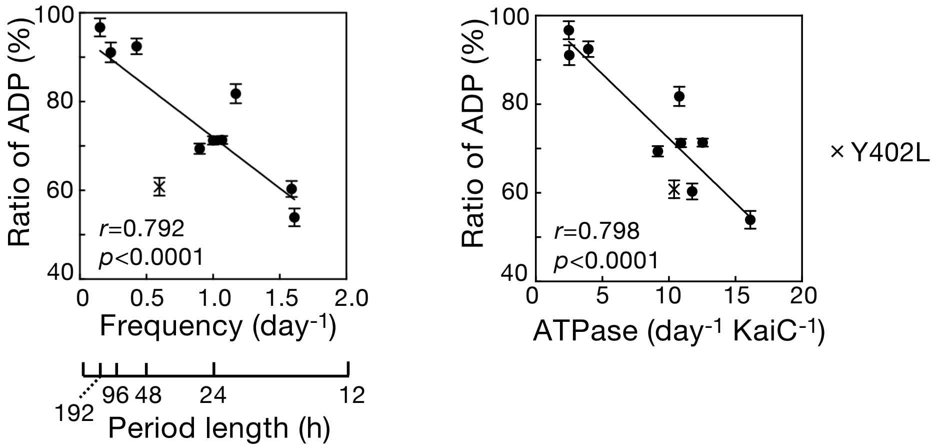


**Fig. S3. Steady-state nucleotide occupancy of WT KaiC and period mutants, including Y402L.**

The fraction of ADP among total KaiC-bound nucleotides is plotted versus the frequency of the in vitro phosphorylation cycle (*left*) or versus KaiC ATPase activity (right). *r*, correlation coefficient; *p*, *P* value. Nucleotide occupancy data are presented as mean ± SD (WT, n = 5; each mutant, n = 3). Frequencies of the in vitro phosphorylation cycle were obtained from (1, 2).

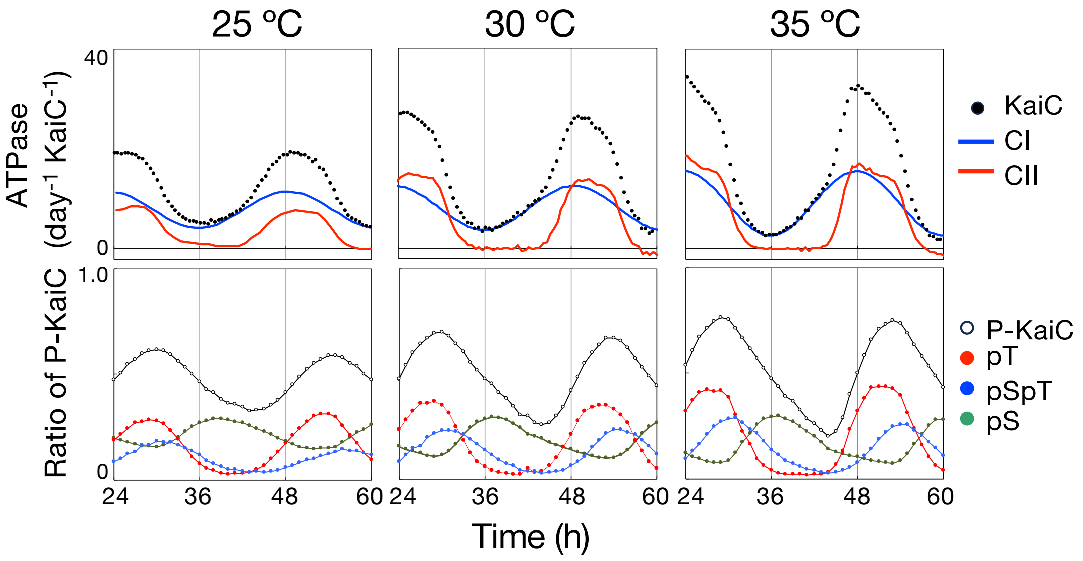


**Fig. S4. Circadian oscillations of KaiC ATPase activity (top) and KaiC phosphorylation (bottom) in an in vitro reconstituted KaiABC reaction at 25°C, 30°C, and 35°C.**

Black circles indicate measured KaiC ATPase activity, whereas blue and red curves denote the inferred CI- and CII-derived ATPase components, respectively. ATPase data are shown as means (n = 3 for 25°C and 35°C; n = 8 for 30°C). Phosphorylation-state data are shown as means (n = 9 for 25°C; n = 11 for 30°C; n = 12 for 35°C).

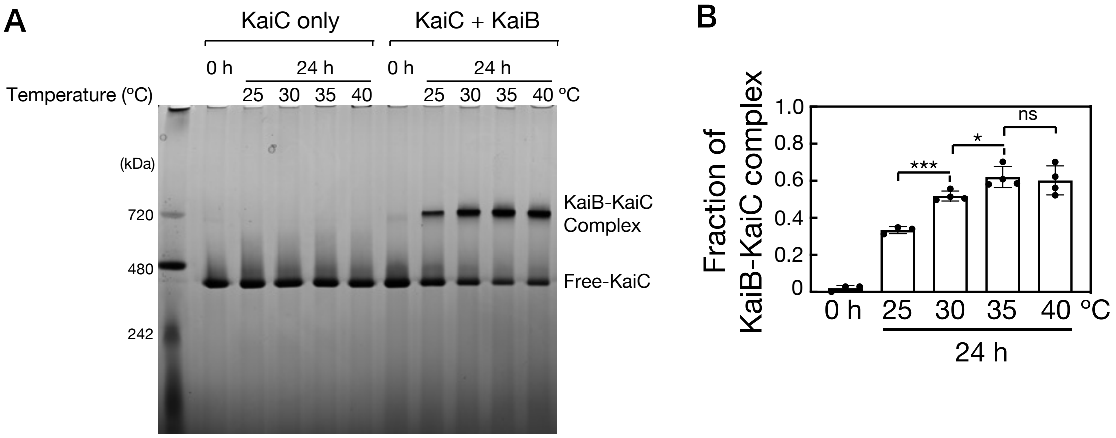


**Fig. S5. Effect of temperature on KaiB–KaiC complex formation.**

(*A*) KaiC was incubated with KaiB for 24 h at 25, 30, 35, or 40°C, and reaction mixtures were analyzed by modified BN-PAGE (see Methods). The upper and lower bands correspond to the KaiB–KaiC complex and free KaiC hexamers, respectively. (*B*) The fraction of KaiB–KaiC complex was calculated as the band intensity of the complex divided by total KaiC signal (complex + free hexamer) and plotted. Statistics were assessed by unpaired *t* tests with Welch's correction; significant differences are indicated as ns (not significant), * ($P < 0.05$) and ***($P < 0.0005$). Data are mean ± SD (n = 3 for 0 h and 25°C; n = 4 for all other temperatures).

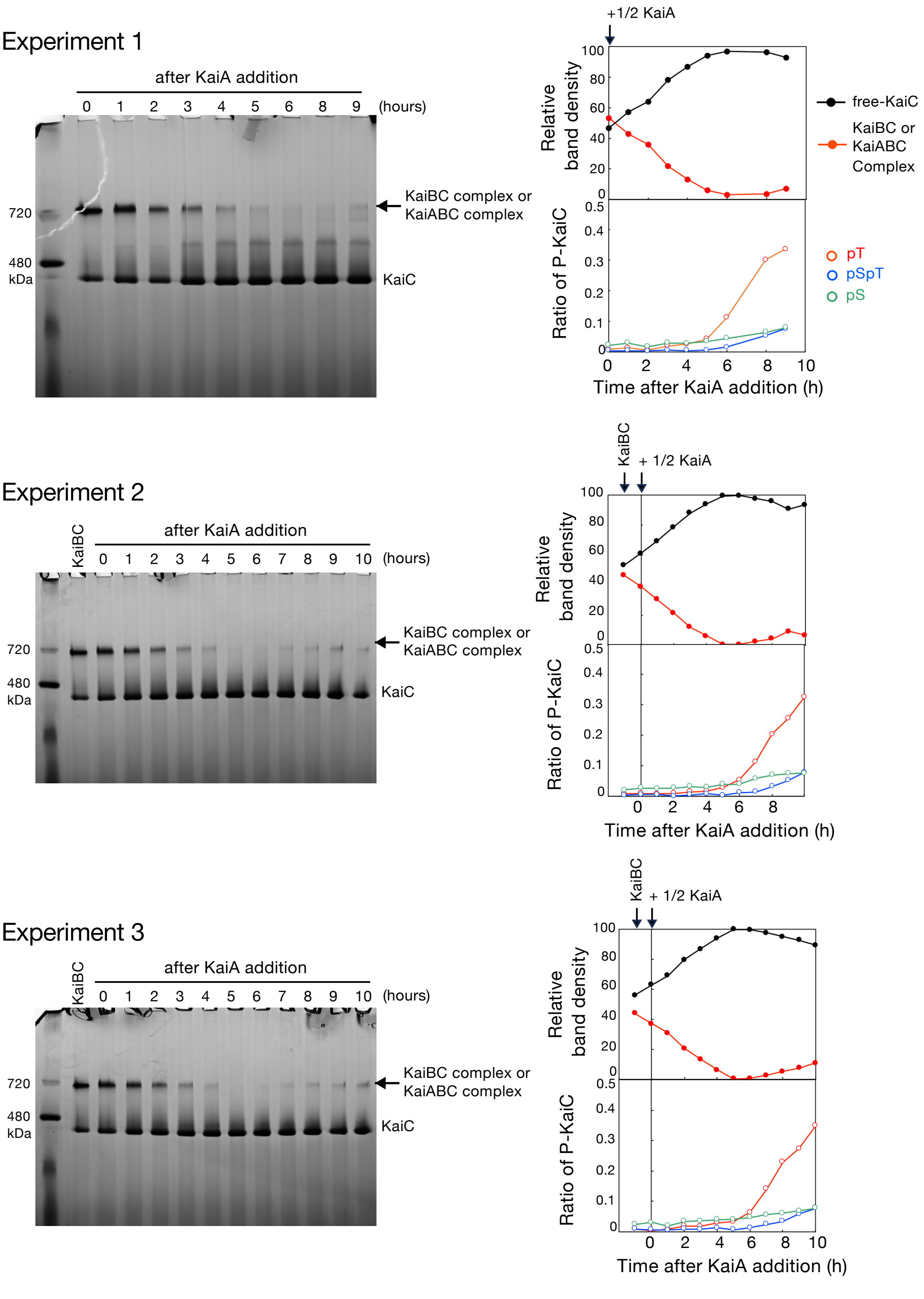


**Fig. S6. Dynamics of KaiB–KaiC complex assembly preceding KaiC phosphorylation.**

Dephosphorylated KaiC was incubated with KaiB for 24 h at 30°C, after which KaiA (0.6 μM) was added. Aliquots were collected at the indicated times and analyzed by modified BN-PAGE or SDS-PAGE. BN-PAGE bands corresponding to free KaiC hexamers and KaiB–KaiC or KaiA–KaiB–KaiC complexes were quantified by densitometry, and the fraction of complexed KaiC was plotted as the complex band intensity divided by total KaiC signal (complex + free hexamer).

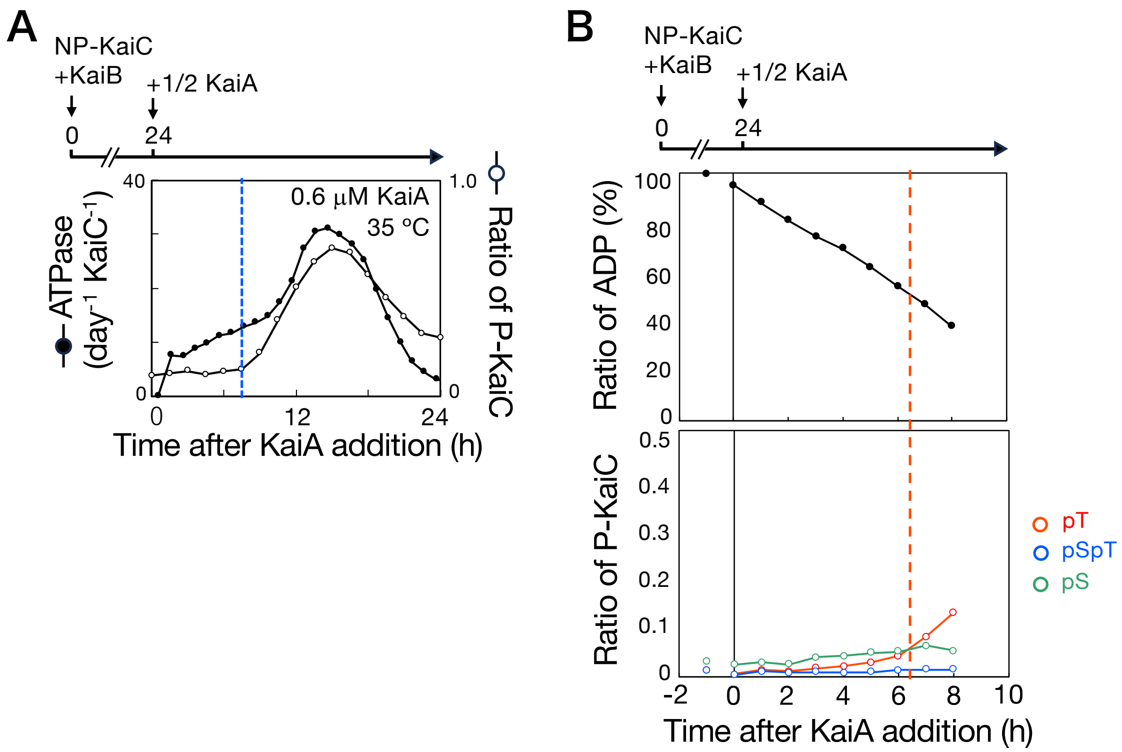


**Fig. S7. KaiA-induced changes in KaiC ATPase activity, nucleotide occupancy, and phosphorylation at 35°C.**

(*A*) Time course of KaiC ATPase activity (filled circles) preceding KaiC phosphorylation (open circles) at 35°C after addition of KaiA (0.6 μM), i.e., half the standard concentration used under oscillatory conditions. The blue dashed line indicates the onset of phosphorylation. (*B*) Time-dependent changes in KaiC nucleotide occupancy (top) and phosphorylation level (bottom) at 35°C following KaiA addition. Dephosphorylated KaiC was preincubated with KaiB for 24 h at 30°C, and KaiA (0.6 μM) was added after 24 h. KaiC ATPase activity, nucleotide occupancy, and phosphorylation were quantified. The red dashed line indicates the time at which phosphorylation begins.

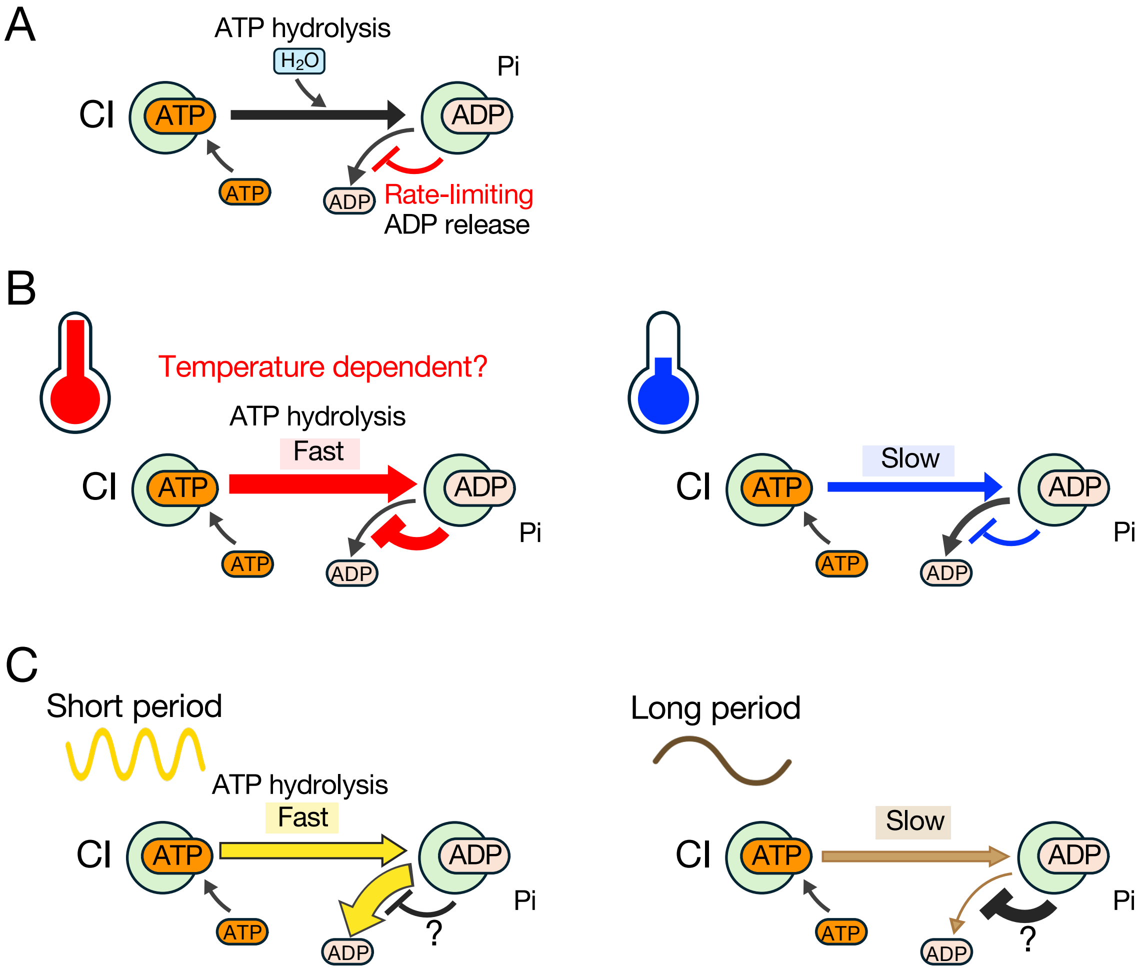


**Fig. S8. Model for negative feedback regulation of KaiC ATPase activity in which ADP release from the CI domain is rate-limiting step.**

(*A*) ADP release from the CI is the rate-limiting step in the KaiC ATPase cycle. (*B*) Temperature compensation. Although ATP hydrolysis in CI is expected to accelerate with temperature by general thermal acceleration of chemical reactions, elevated temperature suppresses ADP release from CI, increasing ADP occupancy and thereby limiting further turnover. (*C*) Period determination. Due to the sequestration of a lytic water molecule in a favorable/unfavorable position in the CI active site, ATP hydrolysis is faster in short-period KaiC mutants and slower in long-period mutants (3). In parallel, short-period mutants release ADP more readily, whereas long-period mutants retain ADP, biasing the equilibrium toward ATP-bound and ADP-bound states, respectively.

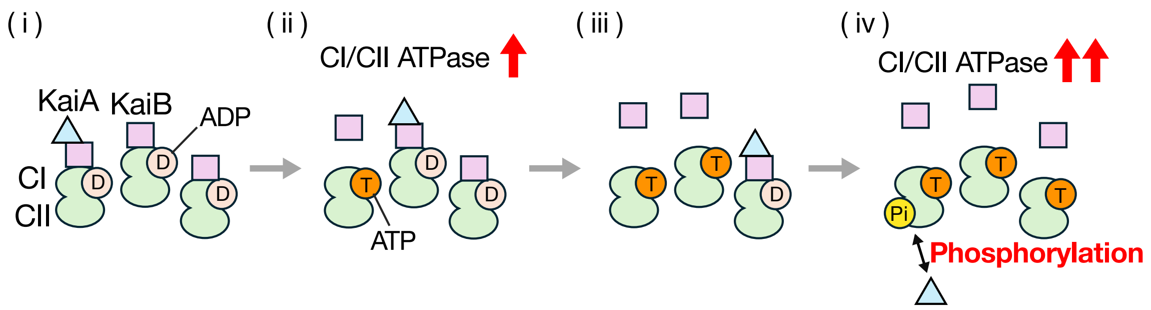


**Fig. S9. CI-domain nucleotide exchange triggers KaiC activation prior to the transition to the CII-domain phosphorylation phase in the KaiABC oscillator at the population level.**

(*i*) KaiA is sequestered to KaiB bound to the ADP-bound CI state. (ii) KaiA-dependent activation of CI/CII ATPase activities promotes KaiB release. Because KaiB suppresses ADP dissociation from CI (Fig. 3), KaiB dissociation is expected to permit CI nucleotide exchange and may thereby further enhance CI ATPase activity. (*iii*) The released KaiA is then captured by another KaiBC complex. (*iv*) As KaiB dissociates and CI becomes predominantly ATP bound state in the reaction mixture, free KaiA triggers the transition to the CII autophosphorylation phase and may further stimulate CI/CII ATPase activity.

**SI References**


1. K. Ito-Miwa, Y. Furuike, S. Akiyama, T. Kondo, Tuning the circadian period of cyanobacteria up to 6.6 days by the single amino acid substitutions in KaiC. *Proc Natl Acad Sci U S A* **117**, 20926-20931 (2020).
2. K. Ito-Miwa, K. Imai, K. Terauchi, T. Kondo, Intrinsic period stability of the cyanobacterial circadian oscillator across in vitro and in vivo conditions. *Proc Natl Acad Sci U S A* **123**, e2526714123 (2026).
3. J. Abe *et al.*, Circadian rhythms. Atomic-scale origins of slowness in the cyanobacterial circadian clock. *Science* **349**, 312-316 (2015).
